# New markers for qPCR detection of the dinoflagellate *Alexandrium catenella* in Chile

**DOI:** 10.1101/2021.10.11.463552

**Authors:** Javiera Espinoza, Kyoko Yarimizu, Satoshi Nagai, Oscar Espinoza-González, Leonardo Guzman, Gonzalo Fuenzalida

## Abstract

*Alexandrium catenella* (Whedon & Kofoid) is a dinoflagellate known as a primary source of paralytic shellfish poisoning in Chile. The distribution range of harmful algal blooms generated by this species has extended during the last decades, and the frequency of these events has increased. In this work, we developed TaqMan markers from Chilean strains that can be used to identify and quantify through qPCR, which can be implemented in monitoring programs for the early detection of this species.

---

Harmful Algal Blooms (HABs) are recurrent events along the Chilean coast, where dinoflagellates species, such as *Alexandrium catenella, Dinophysis acuminata*, *Pseudochattonella verruculosa*, and *Karenia selliformis,* are the culprit for most cases (Díaz et al., 2019; Lagos, 1998; Mardones et al., 2021, 2020). Blooms of *Alexandrium catenella* in Chile have extended their geographical distribution from Southern Patagonia (53°S) to central Chile (36°S) (Guzmán et al., 2002; Mardones et al., 2010), causing massive death of endemic fauna and economic losses to the aquaculture industries (Álvarez et al., 2019; Mardones et al., 2015; Mascareño et al., 2018; Montes et al., 2018). The potent saxitoxin and its derivatives produced by *A. catenella* (Anderson et al., 2012; Krock et al., 2007) can be accumulated in the filter-feeding bivalves at elevated concentrations (Navarro and Contreras, 2010; Pizarro et al., 2018), and humans who consumed the contaminated shellfish can be intoxicated and led to death (García et al., 2004). Implementing molecular tools into algal monitoring is one of the significant innovations because this technology, compared to traditional microscopic detection (Uthermöl, 1958), allows the target species detection with high taxonomic precision even at low concentrations (Anderson et al., 2019; Farrell et al., 2016; Galluzzi et al., 2005: Zhang et al., 2018).

Detection and quantification of molecular markers by quantitative PCR (qPCR) is one of the most attractive strategies for early warning of HAB species to mitigate the impacts of the events. Several prior studies introduced the molecular markers to identify *Alexandrium* species (Antonella & Luca, 2013), convincing that this technique is highly valuable and potentially adaptable for the HAB monitoring systems (Hosoi-Tanabe and Sako, 2005; Kamikawa et al., 2007; Galluzzi et al., 2010; Gao et al., 2015; Garneau et al., 2011; Farrell et al., 2016; Murray et al., 2019; Vandersea et al., 2020). Chile has several on-going HAB monitoring programs (Sandoval et al., 2018; Yarimizu et al., 2020); however, none uses qPCR molecular markers to report the presence of *A. catenella* species. The government institution in charge of protecting public health in Chile considers the *A. catenella* species a hydrobiological plague. The molecular markers that can detect target species with high accuracy can support the current HAB monitoring programs. The purpose of our study was to develop TaqMan-based markers targeting ribosomal RNA genes of the Chilean *A. catenella* strains and to establish a PCR method utilizing these markers to quantify the *in-vitro A. catenella*.

*In-vitro A. catenella* cells (Chilean strain ACENM) and two negative controls, *Pseudochattonella verruculosa* (Hosoi-Tanabe et al., 2007) (Chilean strain CREAN_PV01) and *Chattonella marina* (Hara & Chihara) (Japanese strain CM1), were maintained in sterile flasks (70 ml) containing L1 media at 18◻ with 12-hr light cycle. These culture samples were filtered and processed for DNA extraction by the Chelex-buffer method (Nagai et al., 2012). The extracted gDNA was stable at 5◻ at least for three weeks. The primers and probe were designed from the conserved region by aligning 100 Chilean *A. catenella* sequences (28S rRNA gene) from GenBank, using MUSCLE (Edgar, 2004) implemented in Geneious (Biomatters Ltd): AC-1976F (GTG GGT GGT AAG TTT CAT GCA), AC-2245R (GTG CAA AGG TAA TCA AAT GTC CAC), and probe AC-2026P (FAM-CGC ACA AGT ACC ATG AGG GA-TAMRA). In-silico specificity was checked by Genious software and Primer-BLAST at the NCBI website. The qPCR standard was prepared as follows: The region between the primer set was amplified from the gDNA using KOD-plus-ver.2 (TOYOBO) with the thermal cycle initiation at 94°C for 2 min, 35 cycles of denaturing at 94°C for 10 sec and annealing at 58°C for 30 sec, and extension at 68°C for 40 sec. The product amplicon was verified for 249 bp by agarose gel and purified with a High Pure PCR Product Purification Kit (Roche). The purified amplicon was diluted with DNA/RNA free water to an 8-point standard ranging from 10 to 1×10^−6^ ng/μL. The SYBR qPCR was performed with an 8-point standard to ensure the melt-curve peak at 81 ◻, verifying the target gene amplification. SYBR method used 10 μl sample mix containing 40 nM of each primer and 2×Lightcycler®480 SYBR Green I Master mix (Roche) with thermal cycle of initial enzyme activation at 95 C for 15 min followed by 45 cycles of 95 C for 10sec and 60 C for 30 sec. Six different TaqMan master mix were evaluated by the 8-point standard curve with triplicate samples by the qPCR program; initial incubation at 50 °C for 2 min and 95 °C for 10 min, 45 cycles of 95 °C for 15 s and 60 °C for 1 min. The tested master mix in 25 μL consisted of 2.5 μL of 10 μM primers, and 1 μL of 10 μM probe, 2 μL DNA, 12.5 μL Kapa3G (KAPAbiosystems, KK7251), 0.3 μL of 2.5U/μL Kapa3 plant DNA poly (2.5U/μL) with/out 3 μL of 25 mM MgCl_2_, or the same primer and probe contents with 12.5 μL TaqPath (Applied Biosystems, A30865) with/out MgCl_2_, or those with 12.5 μL Luna universal probe (BioLabs, M3004S) with/out 1.3 μL Clarity JN solution (BioLabs, 12006). To test reproducibility, gDNA was extracted from the two different *A. catenella* culture flasks at two different dates (lot A and B), preparing a 5-point serial dilution from 10 ng/μL gDNA (n=2), which were analyzed by the TaqMan qPCR. The specificity was verified by testing the gDNA (0.1 ng/μL) of *C. marina* and *P. verruculosa* with the TaqMan qPCR. Lastly, the method’s capability to quantify *28S rRNA* gene (LSU) copies from gDNA was tested: *A. catenella* cell count of 30,000, 10,000, and 7,500 estimated by microscope were processed for DNA extraction and assayed by the TaqMan qPCR. The copy number of the target *28S rRNA* gene was determined by the equation, where DNA conc. is SQ value in ng/μL, CF is concentration factor, genome bp is 249 bp, the number of target gene is one based on the assumption of one target *28s rRNA* gene in the region between the primer set.

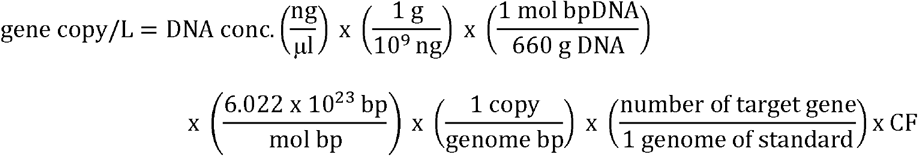

Of six TaqMan master mix evaluated, TaqPath, Luna with JN solution, and Kapa3G with MgCl_2_ provided the best and comparable standard curves. The results obtained with Kapa3G are shown hereafter. The 8-point standard curve showed R^2^=0.997 and efficiency of 102.9 % (Figure 1A, 1B). The standard curve was converted to gene copies (Figure 1C). The reproducibility was confirmed by the comparable TaqMan qPCR results obtained from the two lots of gDNA (Figure 2A, 2B). This result indicated that the method was able to quantify the target gene from the culture gDNA within the range of 10 - 10^−2^ ng/μL (R^2^≧0.99 for both lots). The gDNA at 10^−3^ ng/μL was below the detection limit of the method. The method specificity for *A. catenella* was confirmed as there was no detection of *C. marina,* and the signal from *P. verruculosa* was below the limit of quantitation (Figure 2C, 2D), and the melt peak from these species did not match with that from *A. catenella* (81°C). The target gene copy was average 37,422 per cell, obtained from the *A. catenella* cell count of 30,000, 10,000, and 7,500 assayed by the TaqMan-qPCR. The starting cell count lower than 1,000 showed considerable variability in the resulting copy numbers per cell. When gDNA was extracted from 10,000 cells first and diluted by 10-fold, equivalent to the gDNA quantity from 1,000 cells, the obtained copies per cell were comparable to the average described above. Thus, the cell ratio to a volume at the starting point must be adjusted for the method to obtain reproducible gene copies.

**Figure 1.**
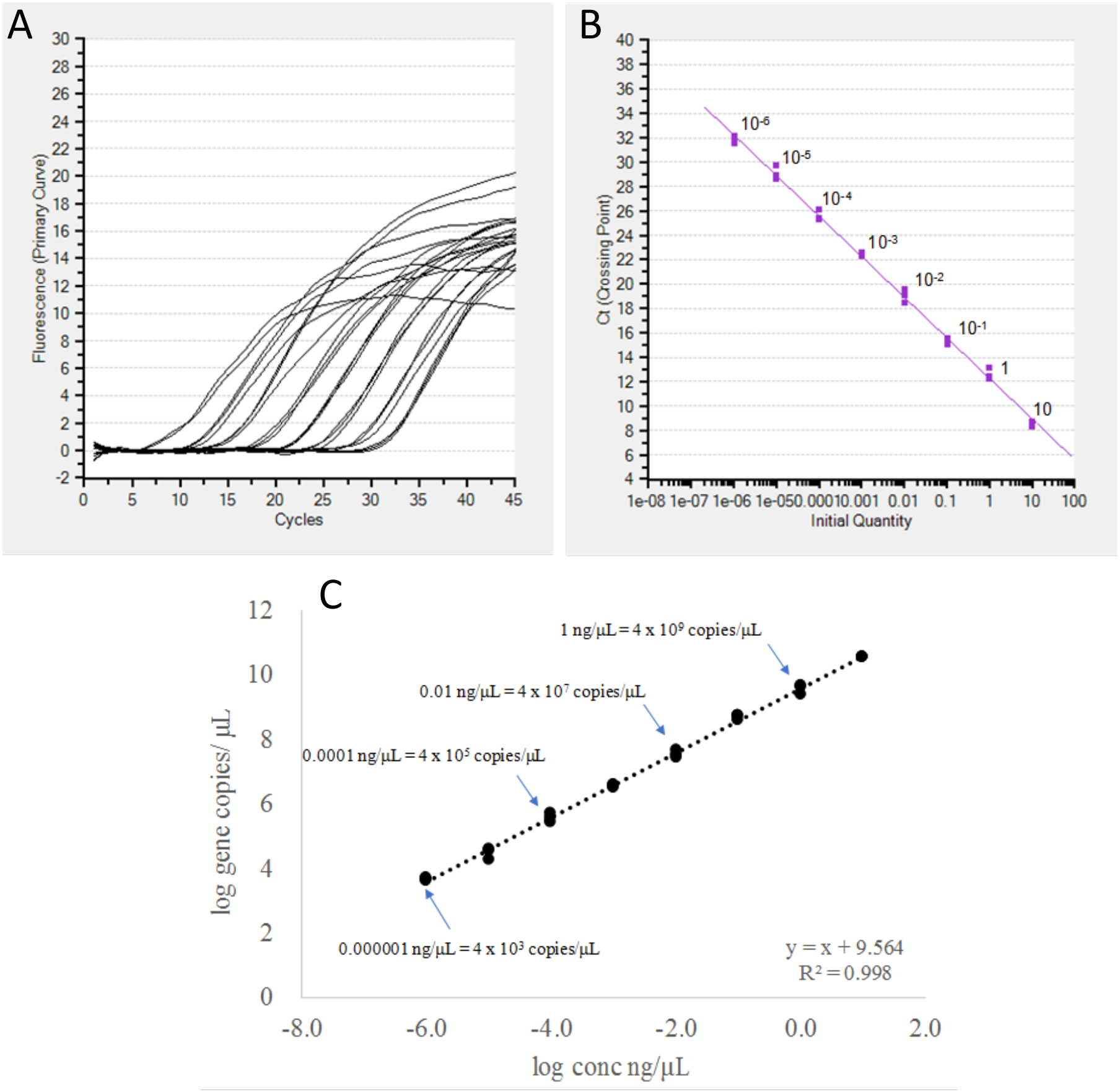
Standard curve of culture-based DNA treated by PCR with the primer AC-1976F and AC-2245R, for the eight-point serial dilution made from 10 ng/μL of standard (n=3) (A and B). Conversion of from concentration to gene copies by calculation based on the assumption that there is one 28s rRNA target gene in the region between primer AC-1976F and AC-2245R (n=3) (C).

**Figure 2.**
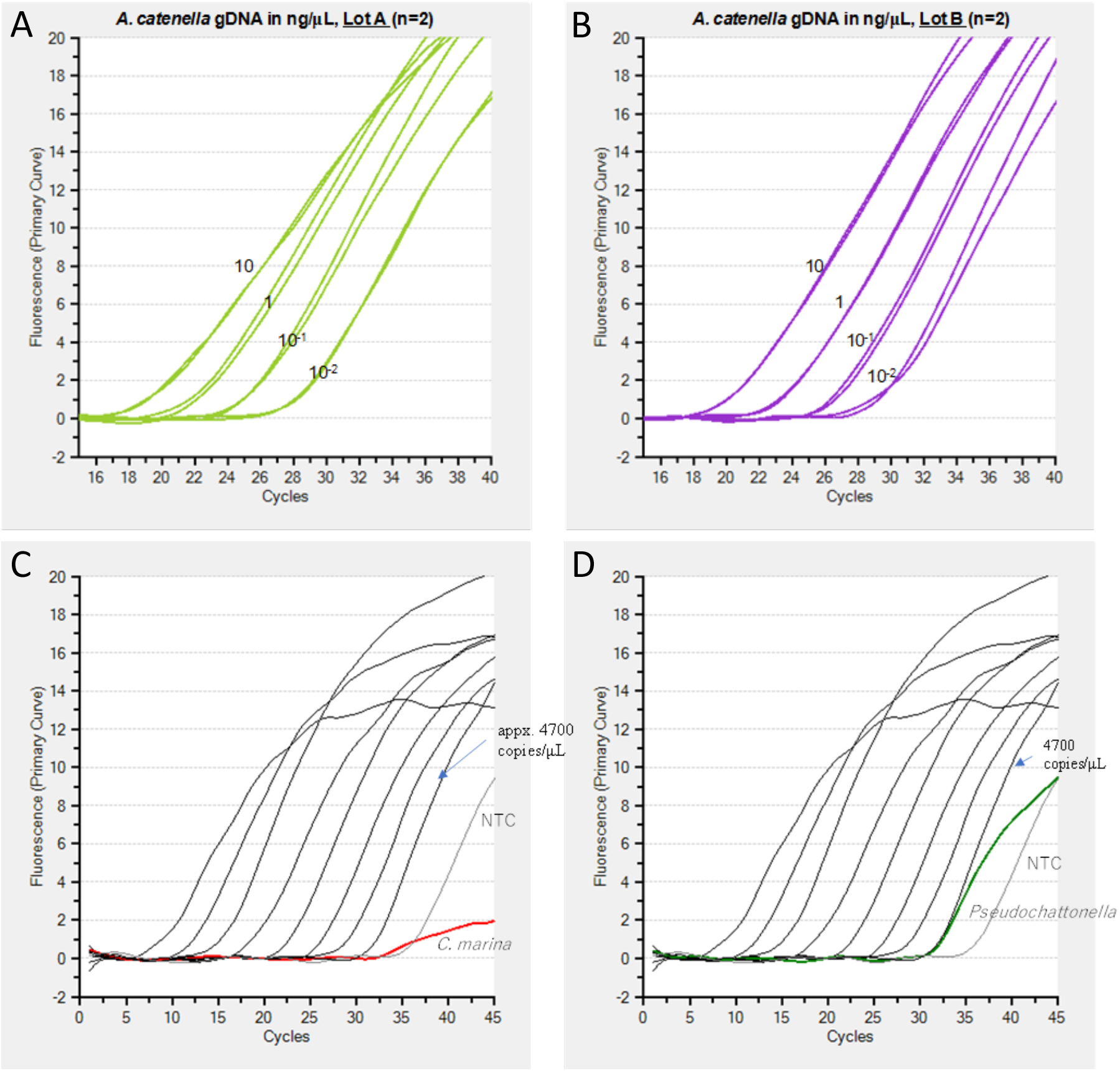
DNA of *A. catenella* from the two different culture flasks tested by the Taq-Man qPCR with Kapa3 master mix. The 5-point serial dilution was made for each DNA based on the 10 ng/ μL of DNA concentration as the stock and the gDNA above 10-2 ng/μL was detected with the linear regression above 0.99 (n=2) (A and B). The gDNA extracted from *C. marina* and *Pseudochattonella* were tested with the *A. catenella* TaqMan-qPCR. There was no detection of *C. marina.* The signal obtained from the *P. verruculosa* was below the lowest standard concentration (C and D).

The present study developed the molecular makers for the Chilean *A. catenella* and established a TaqMan-qPCR method to identify and quantify the species in the cultures. The highlight of this work is that the markers were designed from 100 sequences obtained from Chilean *Alexandrium catenella* strains, aiming to capture the genetic diversity of this species in Chilean coastal monitoring. This is the first marker development targeting the detection of Chilean *A. catenella* strains and has a great potential to serve for early warning tool for Chilean *A. catenella*. The main difference with the work of Murray et al. (2018) is that our makers contain a probe to the qPCR method to increase the specificity, as other *Alexandrium* species have similar sequences. Our preliminary environment sample testing with these makers indicated that the method is quantitative; however, we should keep in mind that the copy number of ribosomal genes is highly variable in *Alexandrium* species (Galluzzi et al., 2010), when the method is applied to environment samples. Overall, the method has a broader potential for field sample surveys to mitigate the HAB impacts. The fisheries equipment transport, ballast water of ships, and the projections under climate change can transport microalgae to spread to broader geographic areas (Griffith and Gobler, 2020; Rodríguez-Villegas et al., 2020), introducing the HABs with new species and further leading to recurrence of the events. The addition of the molecular techniques to the current Chilean HAB monitoring programs can provide comprehensive information and strengthen our knowledge of the phytoplankton dynamics in the regional waters. The method is highly beneficial for early warning of the target HAB species with cell densities, informing the relevant authorities, which can reduce the public health, ecological and economic impacts (Hernandez-Becerril et al., *2*018).

## Acknowledgements

This study was supported by the Instituto de Fomento Pesquero (IFOP) (Grants MR656-114, MR656-123) and the grant (JPMJSA1705) for a study on Science and Technology Research Partnership for Sustainable Development - Monitoring Algae in Chile (SATREPS-MACH).

